# Experience with mating receptivity cues affects sexual behaviour of male guppies, but not their strength of preference towards receptive females

**DOI:** 10.1101/2023.02.07.527553

**Authors:** Versara Goberdhan, Iulia Darolti, Wouter van der Bijl, Judith E. Mank, Alberto Corral-Lopez

## Abstract

Females are traditionally presented as the choosier sex, selecting males based on the quality of their traits. Yet, there is increasing evidence that male mate choice is also important, even in species without male parental care. Social environment and learning are key factors in determining mate preference, and animals are able to use the information they gather from previous experience to potentially increase their odds of obtaining a high-quality mate. We examined how the social environment affects male mate choice in the guppy (*Poecilia reticulata*). We evaluated whether male guppies with previous social experience of female receptivity cues learn to prefer and adapt their behavioural repertoire towards females with higher receptiveness levels, as this represents an optimal use of time and energy and is more likely to result in insemination. For this, we measured sexual preference and behaviour for receptive females in no-choice and dichotomous choice tests using guppy males experienced or naïve to female receptivity cues. Experience with receptivity cues did not change the strength of preference towards receptive females. However, male guppies that had previous experience with female receptivity cues adapted their mating tactic compared to naïve males. The change in mating tactics but lack of preference towards receptive females shows that the influence of social learning is present but might be weaker than predicted in this species. Furthermore, these results provide further support to studies of female mate choice suggesting mating status is not a key factor driving the strength of sexual preferences in natural populations.

## Introduction

Mate choice is a fundamental evolutionary process, as it influences which individuals successfully pass their genes to the next generation (Darwin, 1871; Andersson, 1996). While females historically have been studied as the choosy sex, there is compelling evidence of male mate choice, suggesting that males can be choosy even in species without male parental care (Edward & Chapman, 2011; Herdman et al., 2004; Rosenqvist, 1990; Werner & Lotem, 2003; Wong & Jennions, 2003).

Previous experience can play an important role in forming mating preferences. The social environment can create opportunities for animals to collect information about their surroundings. Animals regularly use this information to adapt their behaviour and efficiently obtain and compete for mates, as well of for other resources that lead them to increase their fitness (Bailey & Moore, 2012; Danchin et al., 2004; Fowler-Finn & Rodríguez, 2012; Valone & Templeton, 2002). In systems in which males perform sexual displays to solicit copulation consent, previous experience can help males to better assess female quality or to avoid soliciting females that signal unwillingness to mate (Akinyemi & Kirk 2019, Dukas 2005, Rather et al., 2022). Similarly, males can adapt their behavioural repertoire or shift to alternative mating tactics based on previous encounters with females (Bailey et al, 2010, Řežucha & Reichard, 2014). Considering the effect of social environment is therefore paramount when evaluating male mate choice patterns.

In fish, male mate choice has been documented in many species (Schlupp, 2018), with males selecting for female traits associated with higher fecundity. For instance, males exhibit preference for larger females with higher fecundity potential in species such as eastern mosquitofish, *Gambusia holbrooki* (Bisazza et al., 1989; Head et al., 2015, Hoysak & Godin, 2007), sailfin mollies, *Poecilia latipinna* (Gumm & Gabor, 2005), and Atlantic mollies, *Poecilia mexicana* (Plath et al., 2006). In fish species with internal fertilization, preference for females showing higher receptiveness levels is theoretically expected, as it is more likely to result in insemination and therefore higher reproductive fitness per unit effort (Bondurianski, 2001, Jordan et al., 2014). However, more research effort is needed to understand pre-copulatory male mate choice in relation to female mating status.

Guppies, fish native to streams of north-eastern South America and the Caribbean, are a traditional model for sexual selection studies. The species is sexually dimorphic, with smaller and colorful males, and have a non-resource based promiscuous mating system (Houde, 1997). Studies of male mate choice in the guppy have revealed that males prefer larger, more fecund females (Corral-Lopez et al., 2018, Dosen and Montgomerie, 2004, Herdman et al., 2004., Jeswiet et al., 2012). Additionally, guppy males adapt their mating tactics and sexual effort in relation to female mating status (Guevara-Fiore et al., 2009; 2010). Studies in guppies have likewise been crucial for our understanding of the role of social environment in sexual selection, and have shown that females shift their preferences for male color traits depending on early social experiences (Rosenqvist & Houde, 1997, Macario et al., 2017). Similarly, rearing conditions and previous success in mating affect subsequent male mating behavioural tactics (Guevara-Fiore, 2012; Guevara-Fiore & Endler, 2018).

While the role the social environment in sexual selection and male mate choice for specific female traits have been studied in guppies, this system provides the additional opportunity to understand how prior social experiences can affect male preference for female mating status. Here we study whether previous social experience with female receptivity cues affect behaviour and strength of preference of male guppies in relation to female mating status. To empirically test this, we experimentally manipulated the social environment of male guppies and quantified their preference and behavioral repertoire in the presence of receptive and non-receptive females. To avoid potential biases introduced by choice of experimental design paradigm (Dougherty & Shuker, 2015), we assessed the role of previous experience in guppy male choice for receptive females using a combination of dichotomous choice and no choice tests. Given theoretical expectations of fitness maximization, we predicted that male guppies would use social learning to shift their preference levels to favour receptive females. Similarly, we predicted that the social environment would affect how males adapt their behavioural repertoire depending on whether they encounter receptive or non-receptive females.

## Methods

### Study system

All guppies used in this experiment originated from a laboratory-adapted stock population, originally collected from the high predation region of the Quaré River (Trinidad & Tobago). Aquaria contained gravel, water filters, and aquatic plants, and all experiments were approved by institutional animal ethics protocols. Fish were raised at a water temperature of 25 °C with a 12:12 light:dark schedule, and fed a daily diet of flake food (Hikari Fancy Food) and live *Artemia* brine shrimp.

We collected newborn guppies from a stock aquarium and held them in nursery aquaria until they could be accurately sexed by the development of a gonopodium, a modified anal fin (Houde, 1997; Liley, 1966), at which point we removed males and held them in male-specific aquaria in groups of seven individuals. During this time, males were not allowed to visualize any females. Once males reached sexual maturity, as evidenced by the development of male colouration, we randomly allocated them into two experimental treatments, experienced and naïve. To ensure all males had similar age and social experience when tested in behavioural experiments and due to logistic reasons, we performed the experiment in two batches that account for half of the individuals each.

To study the preference and behaviour of naïve and experienced males towards females with different mating status, we exposed them to receptive and non-receptive females in dichotomous and no-choice tests (see details below). Sexual receptiveness towards males strongly correlates with the female guppy reproductive cycle. Levels of female receptiveness are highest following parturition of live offspring and for a period of approximately three days in which new ova are commonly fertilized. Receptiveness levels decrease linearly for the following days until they reach minimum levels approximately ten days post-parturition and are maintained in minimum levels until parturition of a new clutch of offspring (approximately 28 days; Liley, 1966, Houde, 1997). Receptiveness in virgin females presents a similar pattern during first reproductive cycle (Houde, 1997). Following theoretical expectations of receptiveness levels towards males and methods in Guevara-Fiore et al. (2010), we housed small groups of virgin females with males in a 1:1 ratio and used them in behavioural tests the following day (receptive females) or 14 days after (non-receptive females).

### Dichotomous choice preference tests

To assess potential differences in preference for receptive females between naïve and experienced males, we measured time associating with receptive and non-receptive females in dichotomous choice tests. We performed two dichotomous choice tests, with an intervening 45 days treatment exposure to females for experienced but not for naïve males. This testing protocol allowed us to determine whether experienced males acquire information about receptivity during their extended exposure to females.

First, to measure baseline preference for receptive females (pre-treatment test), we performed an initial dichotomous choice test on 62 reproductively mature males of similar age (approximately four months old). We photographed each male after behavioural testing using a Canon EOS Rebel T7i camera in a small glass aquarium (5 x 5 x 5 cm) with white walls and a scale for sizing. Camera colour calibration was performed daily with a Calibrite ColorChecker (X-Rite Inc.). Next, we transferred males to treatment 10L aquaria for a 45-day period. Specifically, we transferred each male tested in the pre-treatment test to a separate tank, with half of the males placed with two other virgin males, two virgin females and two non-virgin females of similar age (experienced condition), and the other half placed with two other virgin males of similar age (naïve condition). Naïve males had restricted visual access to tanks with females.

We used photographs to identify males from the pre-treatment tests following the 45 days of experimental treatment, and transferred them in a 3L aquarium three days prior to a second dichotomous choice test (post-treatment test) to allow for sperm replenishment. This avoids biases in preference measurements due to lack in motivation (Pilastro et al., 2002). While biases due to motivation were only expected for experienced males, naïve males were likewise transferred to 3L aquarium three days prior to their second behavioral test. Males were then presented for dichotomous choice post-treatment tests.

We performed all behavioral tests in a circular arena (diameter = 47 cm) sheltered to prevent disruption. We filmed the arena for 15-minute periods using a OBSBOT webcam (1080P at 30 fps) after a five-minute acclimatization period. For accurate identification of fish with tracking software, we placed them in the experimental arena in 20 second intervals. We placed females first in the arena, randomizing the order of placing receptive and nonreceptive females. Additionally, to account for any olfactory cues, we changed water in the arena between tests. To minimize stress, each fish was netted, placed in a glass bowl and transferred to the testing apparatus. For consistency, the tests were always conducted in the morning for a period of 4-6 hours.

We used idTracker to track the position of males and females in video recordings (Perez-Escudero et al., 2014) and to quantify the distance between the male and female for each video frame. To calculate preference for receptive females, we defined the time that a male associated with each female as the number of frames in the video recording that male was < 4 cm (less than two female guppy average body lengths) to each female. Preference ratio was calculated as:

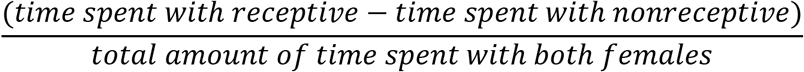

To evaluate differences in preferences for receptive females between experienced and naïve males, we used a Linear Mixed Model with preference ratio as the dependent variable, and the time of testing, experience to mating receptivity cues and the interaction of these as fixed factors. We included the experimental batch as a random factor in the model. Given singularity issues caused by low variance in batch effects, we performed an analogous linear model including batch as a fixed effect. Significance tests were computed using a Wald t-distribution with Kenward-Roger approximation using the parameters package (Lüdecke et al., 2020). All analyses were performed in R (v. 4.1.3; R Core Team, 2022).

To assess potential differences in morphology or colouration patterns between fish used across treatments, we extracted measurements from male photographs taken right after pre-treatment tests. We quantified the number of pixels with carotenoid colouration, black melanic colouration, body size (fish standard length) and tail size in the photographs using ImageJ (Schneider et al., 2012). We used a linear model with each measurement as the dependent variable and social treatments as fixed effect in R (v. 4.1.3; R Core Team, 2022).

### No choice preference tests

To assess potential differences in preference for receptive females, as well as differences in the sexual behaviour repertoire of naïve and experienced males toward receptive females, we performed no-choice tests with receptive and non-receptive females on 122 males from the 31 experienced and 31 naïve aquaria, described above. The males not used in the dichotomous choice tests were presented to either a receptive or nonreceptive female, with 62 experienced males (n=31 with nonreceptive females and n= 31 with receptive females) and 60 naïve males (n=30 with nonreceptive females and n= 30 with receptive females). We excluded two naïve males due to uncertainty with fish labelling. All males were removed from treatment aquaria three days prior to experimentation for sperm replenishment, and to avoid biases in preference measurements due to lack in motivation (Pilastro et al., 2002).

Tests were conducted in a similar fashion to the dichotomous choice tests except only one female was present for each test, alongside a male, with the female placed first in the testing apparatus. Additionally, to account for any olfactory cues, we changed the water between each test. In order to minimize stress, each fish was netted, placed in a glass bowl and transferred to the testing apparatus. For consistency the tests were always conducted in the morning for 4 – 8 hours.

A single observer scored male sexual behaviour in video recordings in a random order and quantified the following behaviours as defined in Liley (1966): i) number of sigmoid displays, every time a male positioned himself in front of the female with an S-shaped posture soliciting copulation; ii) number of sneak attempts, unsolicited attempts of inseminating a female from behind by thrusting his gonopodium at the female’s urogenital pore. We also calculated latency to initial sexual behaviour. Following the procedure described for dichotomous choice tests, we used idTracker software (Perez-Escudero et al., 2014), to calculate the distance between the male and female for each frame in the video recordings and extracted the time spent following a female for each trial (number of frames < 4 cm).

To compare the number of sigmoid displays in experienced and naïve males, we fit a statistical model using a Poisson distribution and a logit link function for the conditional mean in the package glmmTMB (Brooks et al., 2017). We used the mating status of the female, experience to mating receptivity cues and the interaction of these as fixed factors. We included the number of tank and experimental batch as random factors in the model. For sneak attempts and latency to first sexual behaviour, we used analogous models including a zero inflation linear predictor. For time following the female, we used an analogous structure in a Linear Mixed Model fitted with lmer package (Bates et al., 2007). We evaluated the adequacy of our fitted models using scaled-residuals quantile-quantile plots, residual versus predicted values plots and a zero-inflation test in the DHARMa package (Hartig, 2018). We processed the parameters of our statistical models using Wald tests obtained via the parameters package (Lüdecke et al., 2020). We obtained post-hoc comparisons of the male response between female receptivity levels at pre-treatment and post-treatment time of testing in the previous models using the emmeans package with the tukey-adjustment method for multiple comparisons (Lenth et al., 2019). All analyses were performed in R (v. 4.1.3; R Core Team, 2022).

## Results

### Male colour and morphology analyses

Average proportion of orange or black coloration did not differ between males used for experienced or naïve treatments (mean ±SE; orange coloration: naïve males 7.33 ±3.47, experienced males 7.14 ±2.47, F_df=1_ = 121, p = 0.729; black coloration: naïve males 2.27 ±1.02, experienced males 2.20 ±0.86, F_df=1_ = 0.154, p = 0.695; Supplementary Figure 1). Similarly, there was no significant overall difference between naïve and experienced males in morphological traits (body size: naïve males 1.55 ±0.42, experienced males 1.57 ±0.42, F_df=1_ = 0.00, p = 0.99; tail size: naïve males 0.51 ±0.08, experienced males 0.48 ±0.09, F_df=1_ = 0.75, p = 0.38; Supplementary Figure 1).

### Dichotomous Choice Preference Tests

Neither experienced nor naïve males changed their strength of preference towards receptive females after 45 days of treatment (time of testing: estimate_pre-treatment_: −0.01 ±0.04, t = −0.37, p = 0.72; Figure 1; Table1). In addition, we found no overall differences in the strength of preference for receptive females between experienced and naïve males, or in the rate of change in preference between experienced and naïve males following the 45 days of treatment (male social treatment: estimate_experienced_: 0.01 ±0.04, t = 0.45, p = 0.65; male social treatment x time of testing: estimate_pre-treatment x experienced_: 0.04 ±0.06, t = 0.72, p = 0.48; Figure 1; Table 1).

**Figure 1.**
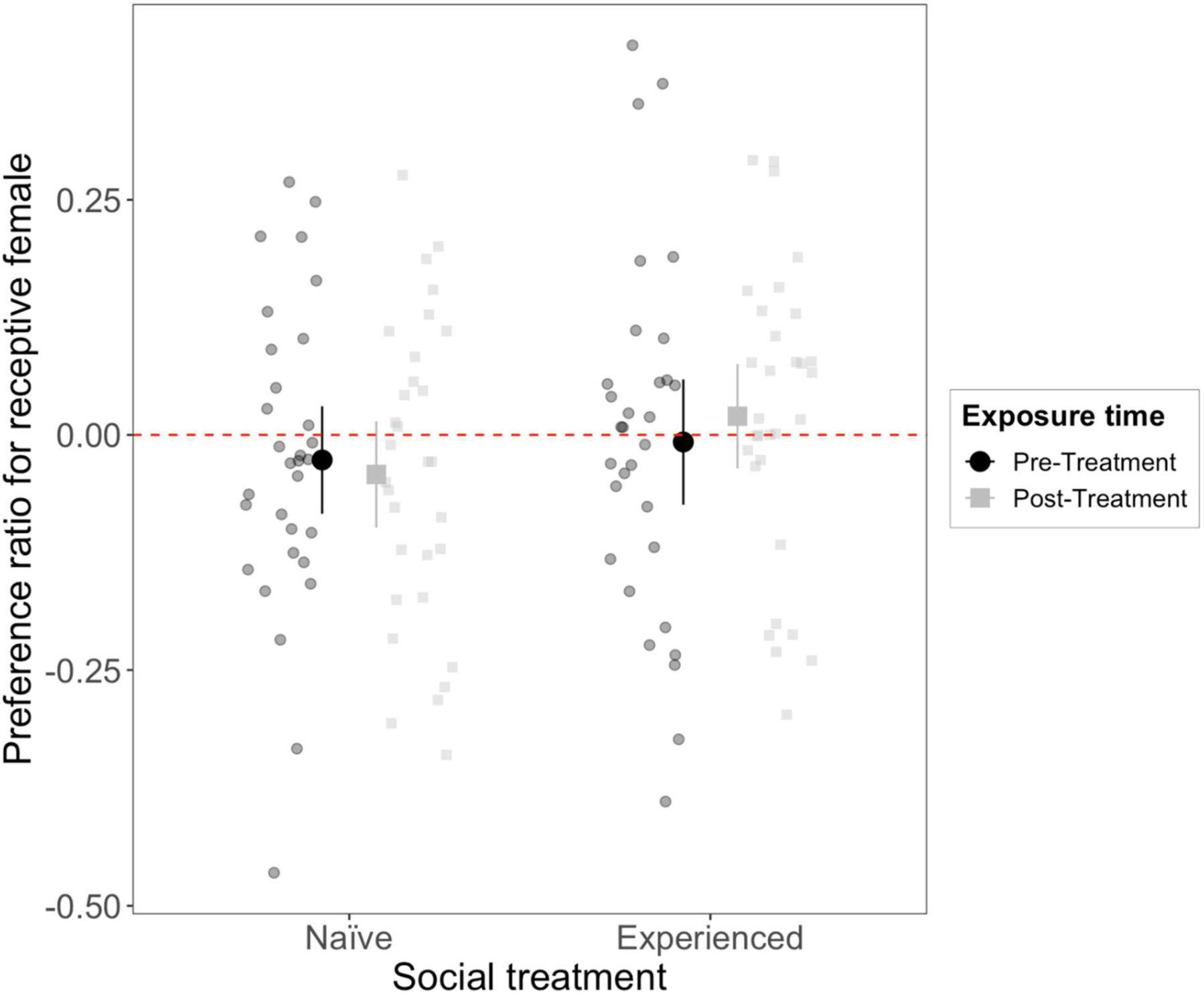
The effect of previous experience with female receptivity cues in guppy male preference for receptive females. Preference ratios were calculated as total time spent associating with receptive females by the total time associating with receptive and non-receptive females in dichotomous choice tests. Tests were performed to males naïve and experienced with female receptivity cues before (pre-treatment, black circles) and after (post-treatment, gray squares) a 45-day treatment in their respective experimental condition. Larger circles and squares indicate average preference ratio for each treatment and time of testing with 95% CI bars. We found no differences in the amount spent with receptive or nonreceptive females between naïve and experienced males in pre-treatment or post-treatment tests (see Table 1).

**Table 1.**
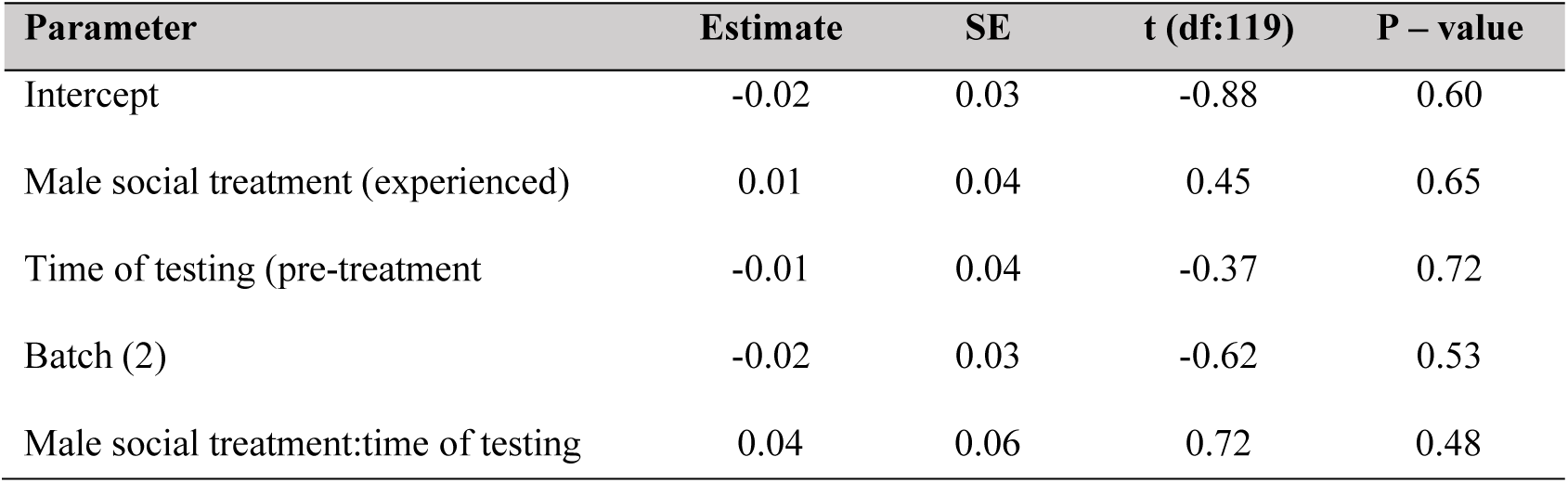
Results for a Linear Mixed Model comparing preference for receptive females performed to male guppies naïve and experienced with female receptivity cues before and after a 45-day treatment in their respective experimental condition.

| Parameter | Estimate | SE | t (df:119) | P – value |
| --- | --- | --- | --- | --- |
| Intercept | -0.02 | 0.03 | -0.88 | 0.60 |
| Male social treatment (experienced) | 0.01 | 0.04 | 0.45 | 0.65 |
| Time of testing (pre-treatment | -0.01 | 0.04 | -0.37 | 0.72 |
| Batch (2) | -0.02 | 0.03 | -0.62 | 0.53 |
| Male social treatment:time of testing | 0.04 | 0.06 | 0.72 | 0.48 |

### No Choice Preference Tests

There was no significant difference between naïve and experienced males in their levels of display behaviour or the number of displays that were performed towards receptive versus non-receptive females (Figure 2a; Table 2). However, we found that, unlike naïve males, experienced males significantly increased the number of displays towards receptive females compared to non-receptive females (female status x social treatment: estimate_receptive x experienced_: 0.26 ±0.08, t = 3.01, p = 0.002; Figure 2a; Table 2).

**Figure 2.**
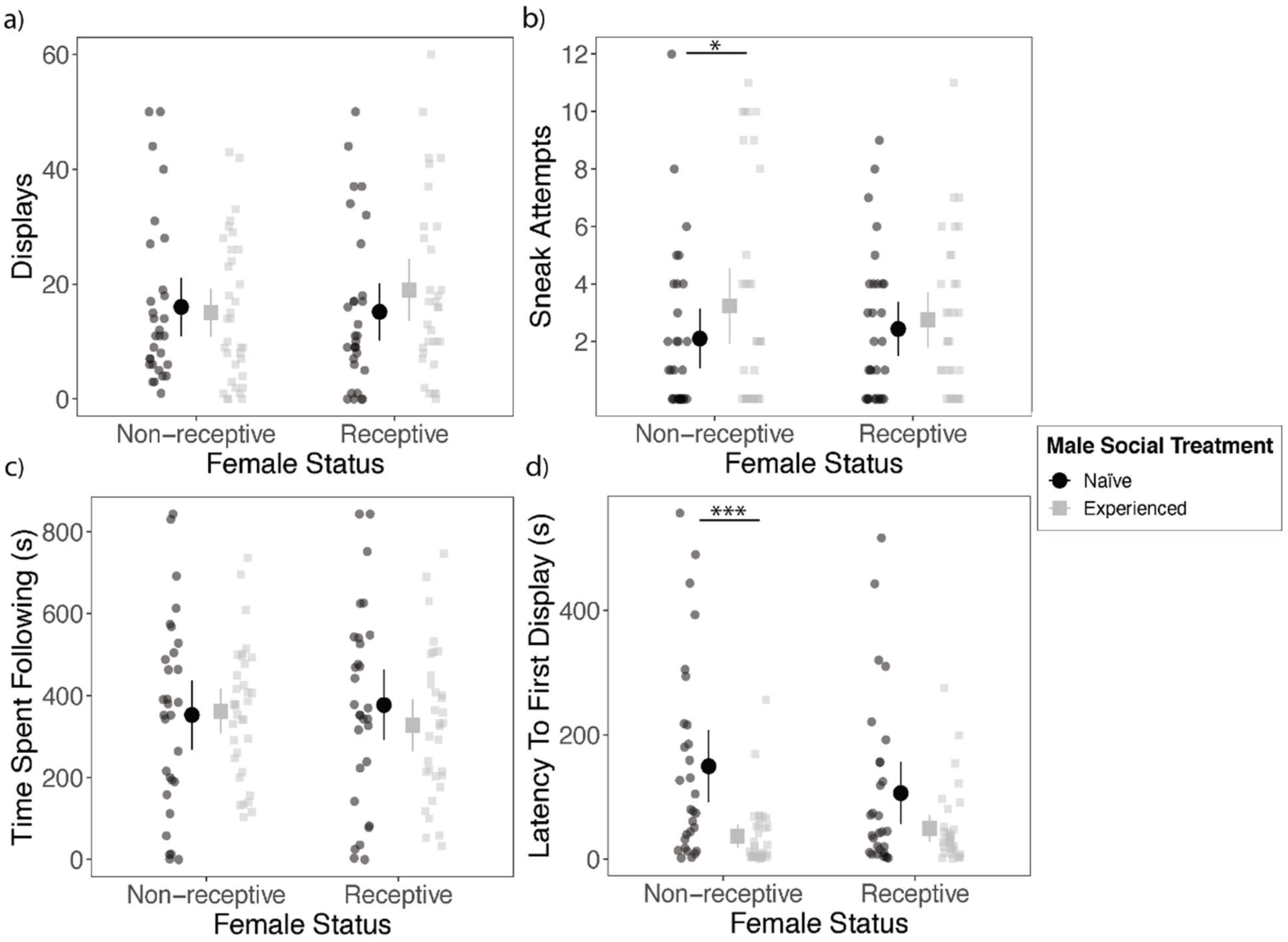
The effect of previous experience with female receptivity cues in male sexual behaviour. Quantification of (a) the number of sigmoid displays, (b) number of sneak attempts, (c) total time spent following females, and (d) latency to first sexual behaviour performed towards non-receptive and receptive females in guppy males naïve (black circles) and experienced (gray squares) with female receptivity cues. Larger circles and squares indicate average values of each behaviours with 95% CI bars. For number of displays, experienced males significantly increased the number of displays towards receptive females compared to non-receptive females (female status x social treatment: p = 0.002; see Table 2). Stars indicate significance in post-hoc comparisons of the male response between female receptivity levels at pre-treatment and post-treatment time of testing (p < 0.001 ***, p < 0.05 *).

**Table 2.** Statistical tests for models comparing potential differences in behaviour in no choice tests with non-receptive and receptive females performed to male guppies naïve and experienced with female receptivity cues. Parameters with significant differences in bold.

| Behaviour | Parameter | Estimate | SE | Statistic<sub>df</sub> | P - value |
| --- | --- | --- | --- | --- | --- |
| *Display* | Intercept | 2.57 | 0.20 | z : 12.70 | < **0.001** |
|  | Female status (receptive) | -0.05 | 0.06 | z : -0.85 | 0.39 |
|  | Male social treatment (experienced) | -0.05 | 0.17 | z : -0.28 | 0.77 |
|  | Female status:male social treatment | 0.26 | 0.08 | z : 3.01 | **0.002** |
| *Sneak* | Intercept | 1.15 | 0.17 | z : 6.48 | < **0.001** |
|  | Female status (receptive) | -0.01 | 0.20 | z : -0.07 | 0.94 |
|  | Male social treatment (experienced) | 0.49 | 0.22 | z : 2.19 | **0.028** |
|  | Female status:male social treatment | -0.47 | 0.26 | z : -1.79 | 0.07 |
|  | *Zero Inflation:* Intercept | 0.49 | 0.10 | z : -3.37 | <**0.001** |
| *Latency* | Intercept | 4.27 | 0.26 | z : 16.42 | < **0.001** |
|  | Female status (receptive) | -0.52 | 0.36 | z : -1.42 | 0.15 |
|  | Male social treatment (experienced) | -1.51 | 0.36 | z : -4.17 | < **0.001** |
|  | Female status:male social treatment | 0.99 | 0.51 | z : 1.93 | 0.053 |
|  | *Zero Inflation:* Intercept | 0.00 | 0.00 | z : -0.00 | 0.996 |
| *Time spent following* | Intercept | 351.27 | 44.13 | t<sub>3,7</sub> : 7.95 | **0.002** |
|  | Female status (receptive) | 24.51 | 42.85 | t<sub>61,5</sub> : 0.57 | 0.56 |
|  | Male social treatment (experienced) | 13.70 | 51.52 | t<sub>105,5</sub> : 0.26 | 0.79 |
|  | Female status:male social treatment | -61.06 | 59.30 | t<sub>61,8</sub> : -1.03 | 0.31 |

Experienced males exhibited significantly more sneak attempts than naïve males (male social treatment: estimate_experienced_: 0.49 ±0.22, t = 2.19, p = 0.028; Figure 2b; Table 2), and this difference was due to greater frequency of sneak attempts by experienced males toward non-receptive females. We observed no difference in sneak frequency between experienced and naïve males in tests with receptive females (log_mean ± SE_; non-receptive females: naïve males 1.16 ±0.18, experienced males 1.65 ±0.14, t-ratio_df=119_ = −2.19, p = 0.030; log_mean ±SE_; receptive females: naïve males 1.14 ±0.16, experienced males 1.16 ±0.15, t-ratio_df=119_ = −0.07, p = 0.94; Fig. 2b; Table 2). Additionally, naïve and experienced males showed no significant difference in the time spent following females in the trials, or in overall time spent following receptive and non-receptive females (Figure 2c; Table 2).

Experienced males showed decreased latency to perform sexual behavior compared to naïve males in tests with non-receptive and receptive females (male social treatment: estimate_experienced_: −1.51 ±0.36, z = −4.17, p < 0.001; Figure 2d; Table 2) However, post-hoc tests indicate that this decrease was significant just in tests with non-receptive females (log_mean ±SE_; non-receptive females: naïve males 4.28 ±0.26, experienced males 2.76 ±0.25, t-ratio_df=115_ = 4.17, p < 0.001; mean ±SE; receptive females: naïve males 3.75 ±0.26, experienced males 3.23 ±0.25, t-ratio_df=115_ = 1.44, p = 0.15; Fig. 2d; Table 2).

## Discussion

We used dichotomous choice and no-choice tests to investigate how previous experience with female receptivity cues alter guppy male sexual behaviour and strength of preference for female mating status. Our results showed that males with previous access to female receptivity cues exhibited significantly greater frequency of coercive sexual behaviours and lower latency to first sexual behaviour to non-receptive females than naïve males. In addition, only experienced males significantly increased their number of displays towards receptive females compared to the number of displays performed with non-receptive females. However, previous experience with receptivity cues did not affect the strength of guppy male preference for female mating status.

Previous studies evaluating how female mating status affect male mating behavior showed higher levels of coercive copulation attempts towards non-receptive females and higher levels of sigmoid displays towards receptive females (Guevara-Fiore et al., 2010a;2010b). Our results match these patterns only in males with previous experiences with female receptivity cues, suggesting a key role of social learning driving preferences for high quality females in this species. Our experimental design does not allow to disentangle which mechanism leads to changes in behaviour between experienced and naïve males. Prior experience with females might lead to males better recognizing which mating tactics provide higher success, as previously observed in species such as *Drosphila melanogaster* (Dukas, 2005, Saleem et al., 2014, Balaban-Feld & Valone., 2017), or eastern mosquitofish (Bisazza et al., 1996 but see Iglesias-Carrasco et al., 2019). Alternatively, changes in encounter rates of females and in mating success artificially created by the two social environments used in our experimental setup are known to affect male mating tactics (Cattelan et al., 2016; Devigili et al., 2015; Jordan & Brooks, 2012, Guevara-Fiore & Endler, 2018). Ultimately, the behavioural patterns observed in males with access to female receptivity cues correspond to theoretical predictions of fitness maximization, once accounting for the lower energetic requirements of sneak attempts of sperm insemination in relation to more costly sigmoid displays aiming to engage female with sexual consent (Devigli et al., 2013, Head et al., 2010).

Contrary to our prediction, males with social learning experience of female receptivity cues did not became choosier or increase their preference towards receptive females in dichotomous choice tests. It may be that that our experimental treatment might have changed the perception of naïve and experienced males in future reproductive opportunities, potentially biasing the investment of naïve males in sexual behaviors that we observed in the first sexual encounter of their pre-treatment test (Fischer et al., 2008, Aich et al., 2021). Furthermore, determining the costs of sexual behaviours are challenging in benign lab environments where food is not a limiting factor. Future work incorporating resource limitation will be helpful to determine ecologically-relevant effects of social learning in male mating preferences. Yet, it is important to note that our results are concordant with a recent meta-analysis showing no evidence that mating status is an important factor for preference. Specifically, across species, virgin females are as choosy as mated females across reproductive isolation, inbreeding avoidance, and sexually transmitted disease scenarios (Richardson & Zuk, 2022). Our study here presents a similar finding, as males with no previous mating experience (naïve males) presented similar preferences to males with mating experience (experienced males). Hence, our study focusing on male mate choice on preference for higher quality females is in broad agreement to previous observations on the role of mating status for female preference across species.

Using both dichotomous and no-choice approaches allowed for a broader picture of male preference in the context of social learning. It is possible that mating preferences may be stronger in choice tests compared to no-choice design, as males can select the female that is more likely to result in insemination (Dougherty, 2020). However, there is arguably an increased risk of being rejected by the only potential mate in a no-choice tests, and this could make males more careful in tuning their mate strategy to female receptivity cues (Dougherty & Shuker, 2015). Our tests using these two complementary experimental paradigms are concordant in that we did not observe changes in overall preference for receptive females or differences in overall sexual behaviour levels based on female receptivity status. However, our observations of changes in rates of sexual display and coercive copulations depending on female mating status suggest that future mate choice studies should incorporate both methodologies.

The changes we observe in male guppies in behavioural repertoire and latency towards females based on previous experience with female receptivity cues add to the evidence suggesting that social environment and learning from previous experience can affect male sexual behaviour. Overall, our results are consistent with the idea that male guppies use social learning to efficiently tune their mating tactics, soliciting copulation in higher rates to receptive females and performing higher coercive copulation attempts towards non-receptive females.

## Acknowledgements

We thank Wouter van der Bijl for help with photography and fish care, as well as Lydia Fong, Yuying Lin and Jacelyn Shu for help with fish care.

## Author contributions

J.E.M. and AC-L. conceptualized the study. V.G. and I.D. performed research. A.C-L and V.G. performed formal analyses and visualization. J.E.M. contributed to funding acquisition of the project. A.C-L., J.E.M., and V.G. wrote the original draft. All authors contributed to the final version of the manuscript.

## Supplementary Materials

**Table S1.** Results of linear model comparing colouration and morphological traits in male guppies used for social experience experimental treatments

| Trait | Sum Sq | Mean Sq | F value (df = 1) | P value |
| --- | --- | --- | --- | --- |
| Orange Colouration | 1.1 | 1.09 | 0.12 | 0.72 |
| Black Colouration | 0.14 | 0.13 | 0.15 | 0.69 |
| Standard Length (cm) | 0.00 | 0.00 | 0.00 | 0.93 |
| Tail Length (cm) | 0.012 | 0.01 | 3.08 | 0.08 |

**Figure. S1.**
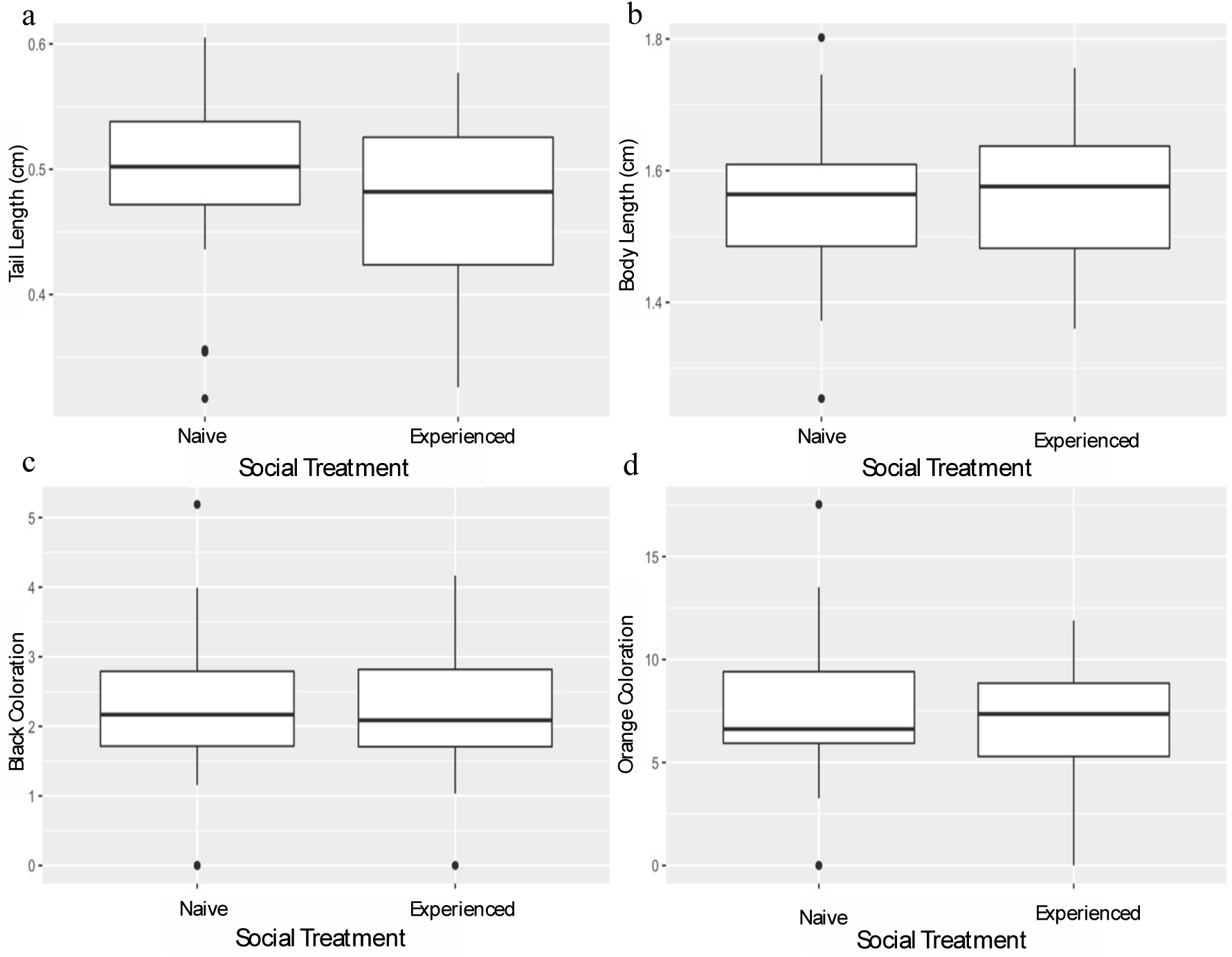
Body morphology and colouration in experienced and naïve males. No significant differences were found between randomly assigned males to naïve (n = 31) and experienced (n = 31) treatments for (a) tail length, (b) body length, (c) proportion of black, or (d) proportion of orange. For all boxplots, horizontal lines indicate medians, boxes indicate the interquartile range, and whiskers indicate all points within 1.5 times the interquartile range.

